# PlaScope: a targeted approach to assess the plasmidome of *Escherichia coli* strains

**DOI:** 10.1101/334805

**Authors:** G. Royer, J.W. Decousser, C. Branger, C. Médigue, E. Denamur, D. Vallenet

**Affiliations:** Assistance Publique-Hôpitaux de Paris, Hôpital Henri Mondor, Université Paris Est Créteil, Département de Microbiologie, F-94000 Créteil, France; Université Paris Diderot, INSERM, IAME, UMR 1137, Sorbonne Paris Cité, F-75018 Paris; LABGeM, Génomique Métabolique, Genoscope, Institut François Jacob, CEA, CNRS, Univ Evry, Université Paris-Saclay, 91057 Evry, France; Assistance Publique-Hôpitaux de Paris, Hôpital Henri Mondor, Université Paris Est Créteil, Département de Microbiologie, Next-Generation Sequencing Platform, F-94000 Créteil, France; Assistance Publique-Hôpitaux de Paris, Hôpital Bichat, Laboratoire de Génétique Moléculaire, F-75018 Paris, France

## Abstract

Plasmid prediction may be of great interest when studying bacteria such as *Enterobacteriaceae*. Indeed many resistance and virulence genes are located on such replicons and can have major impact in terms of pathogenicity and spreading capacities. Beyond strains outbreak, plasmids outbreaks have been reported especially for some extended-spectrum beta-lactamase or carbapenemase producing *Enterobacteriaceae*. Several tools are now available to explore the “plasmidome” from whole-genome sequence data, with many interesting and various approaches. However recent benchmarks have highlighted that none of them succeed to combine high sensitivity and specificity. With this in mind we developed PlaScope, a targeted approach to recover plasmidic sequences in *Escherichia coli*. Based on Centrifuge, a metagenomic classifier, and a custom database containing complete sequences of chromosomes and plasmids from various curated databases, it performs a classification of contigs from an assembly according to their predicted location. Compared to other plasmid classifiers, Plasflow and cBar, it achieves better recall (0.87), specificity (0.99), precision (0.96) and accuracy (0.98) on a dataset of 70 genomes containing plasmids. Finally we tested our method on a dataset of *E. coli* strains exhibiting an elevated rate of extended-spectrum beta-lactamase coding gene chromosomal integration, and we were able to identify 20/21 of these events. Moreover virulence genes and operons predicted locations were also in agreement with the literature. Similar approaches could also be developed for other well-characterized bacteria such as *Klebsiella pneumoniae.*

**Data summary:**

1. All the genomes were downloaded from the National Center for Biotechnology Information Sequence Read Archive and Genome database (Supplementary table 1 and 2).
2. The source code of PlaScope is available on Github (https://github.com/GuilhemRoyer/PlaScope).

**Importance:** Plasmid exploration could be of great interest since these replicons are pivotal in the adaptation of bacteria to their environment. They are involved in the exchange of many genes within and between species, with a significant impact on antibiotic resistance and virulence in particular. However, plasmid characterization has been a laborious task for many years, requiring complex conjugation or electroporation manipulations for example. With the advent of whole genome sequencing techniques, access to these sequences is now potentially easier provided that appropriate tools are available. Many softwares have been developed to explore the plasmidome of a large variety of bacteria, but they rarely managed to combine sensitivity and specificity. Here, we focus on a single species, *E. coli*, and we use the many data available to overcome this problem. With our tool called PlaScope, we achieve high performance compared with two other classifiers, Plasflow and cBar, and we demonstrate the utility of such an approach to determine the location of virulence or resistance genes. We think that PlaScope could be very useful in the analysis of specific and well-known bacteria.

## Introduction

Recently, several studies have evaluated the effectiveness of *in silico* plasmid prediction tools (1, 2). In fact, many bioinformatics methods are now available to detect such mobile elements, with different approaches like read coverage analysis, k-mer based classification, replicon detection; some being fully automatized (3-7), others not (8). Some of them achieve high sensitivity: for example, PlasmidSPAdes and cBar enable plasmid recall of 0.82 and 0.76 on a dataset of 42 genomes, respectively (1). On the other side some tools display very high precision, as PlasmidFinder which reaches 100% (1). Unfortunately, neither of them succeeds in finding a good trade-off between sensitivity and specificity, and thus users need to combine different methods to get correct predictions.

Concomitantly more and more sequences are available in public databases, with various level of completeness from large sets of contigs to fully circularized genomes and plasmids. Some people have made an effort to curate these databases and proposed high quality dataset. Carattoli *et al.* and Orlek *et al.*, for example, have published interesting and exhaustive plasmid datasets for *Enterobacteriaceae* (4, 9).

With this in mind, we propose here a method, called PlaScope, to assess the plasmidome of genome assemblies. We took advantage of available genomic data to create a custom database of plasmids and chromosomes, which is used as input of the Centrifuge software, a tool originally developed as a metagenomics classifier (10). We compared it with others plasmid classifiers, cBar and Plasflow, and showed that with our specific knowledge-based approach we were able to recover nearly all plasmids of various *Escherichia coli* strains without compromising on specificity. Finally, the usefulness of our approach is illustrated on a set of *E. coli* whole genomes for which we have sought to identify the location of specific genes involved in virulence or antibiotic resistance.

## Theory and implementation

### Workflow description

PlaScope workflow is illustrated in Fig. 1. First, users have to provide paired end fastq files. Then assembly is run using SPAdes 3.10.1 (11) with the “careful” option to obtain contigs. Subsequently, Centrifuge (10) predicts the location of these contigs thanks to a custom database and sorts sequences into 3 classes: plasmid, chromosome and unclassified. The latter includes sequences shared by both categories (i.e. plasmid and chromosome) and which are therefore indistinguishable, and sequences without any hit. Finally results are sorted based on those three classes and extracted using awk. The complete workflow is available through a unique bash script called PlaScope.sh on github (https://github.com/GuilhemRoyer/PlaScope).

**Figure 1.**
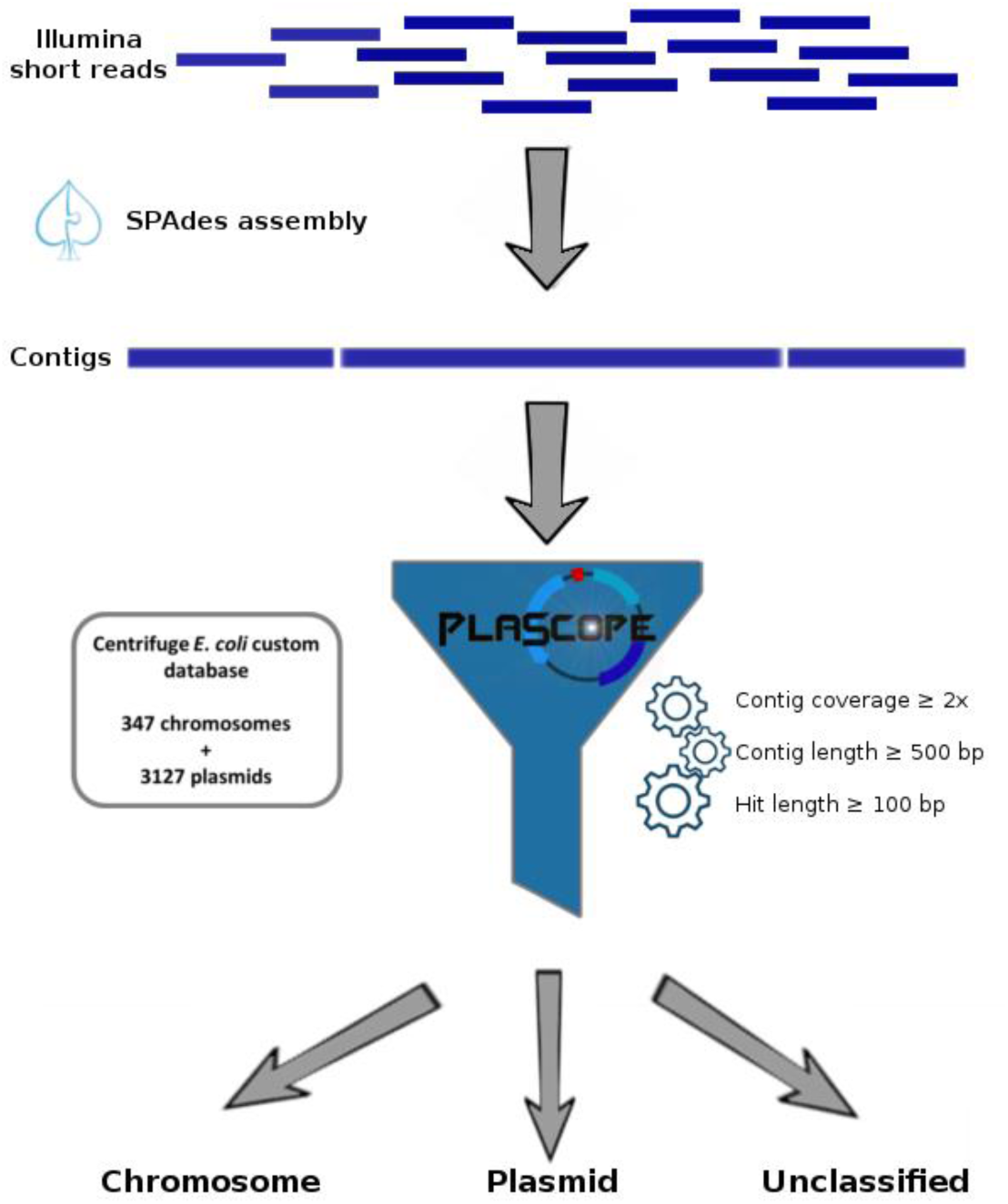
PlaScope workflow. After read assembly using SPAdes, PlaScope classifies contigs into three categories (i.e. chromosome, plasmid, unclassified) based on a custom database of 347 chromosomes of *E. coli* and 3127 plasmids of *Enterobacteriaceae*.

### Centrifuge custom database construction

We gathered all the complete genome sequences (chromosomes and plasmids) of *E. coli* from the NCBI on 10/01/2018. We also added the plasmid sequences that were used to create PlasmidFinder database (4) and those proposed by Orlek *et al.* (9). Finally, we added a very specific dataset containing *E. coli* plasmids involved in antibiotic resistance (www.agence-nationale-recherche.fr/en/anr-funded-project/) (https://www.ebi.ac.uk/ena/data/view/PRJEB24625) (Branger et al., unpublished results). Altogether the database includes 347 chromosome and 3127 plasmid sequences (supplementary table 1 - database is available here https://zenodo.org/record/1245664#.WvXAVaSFNaQ).

**Table 1.** PlaScope, Plasflow and cBar benchmark results on plasmid contigs.

|  | **PlaScope** | **Plasflow** | **cBar** |
| --- | --- | --- | --- |
| **True Positive** | 1123 | 1106 | 954 |
| **True Negative** | 9162 | 6231 | 5570 |
| **False Positive** | 52 | 2983 | 3644 |
| **False Negative** | 173 | 190 | 342 |
| **Recall** | 0.87 | 0.85 | 0.74 |
| **Precision** | 0.96 | 0.27 | 0.21 |
| **Specificity** | 0.99 | 0.68 | 0.6 |
| **Accuracy** | 0.98 | 0.7 | 0.62 |
| **F1 score** | 0.91 | 0.41 | 0.32 |

Then, we pooled separately plasmid and chromosome sequences to create a custom database for Centrifuge 1.0.3 (10) with an artificial taxonomy containing only three nodes: “chromosome”, “plasmid”, and “unclassified” (see README on https://github.com/GuilhemRoyer/PlaScope).

### PlaScope classification method

PlaScope classifies contigs as “chromosome”, “plasmid” or “unclassified” with Centrifuge using our custom database (centrifuge -f --threads 2 -x custom_database -U example.fasta -k 1 --report-file summary.txt -S extendedresult.txt), with the option “k” set to 1 in order to get only one taxonomic assignment. Only contigs longer than 500 bp, with a Centrifuge hit longer than 100 bp and with a SPAdes contig coverage higher than 2 are classified as plasmid or chromosome-related.

### Reference dataset for method evaluation

To evaluate our tool, we searched for completely finished genomes of *E. coli* with Illumina reads available on the National Center for Biotechnology Information (NCBI) database. All corresponding chromosome and plasmid sequences and Illumina short reads were downloaded from the NCBI on 10/01/2018, and converted into fastq files with fastq-dump from sra-toolkit (fastq-dump --split-files). For evaluation purpose, these genomes were not included in the centrifuge custom database.

The short reads were assembled with SPAdes 3.10.1 (11) with standard parameter and “careful” option (spades.py --careful -t 8 −1 read_1.fastq.gz −2 read_2.fastq.gz -o output_directory). After assembly, 16S rapid identification was performed on fasta files using ident-16s (12). 12 assemblies which did not contained *Escherichia* 16S or with multiple 16S from various organisms were excluded from the subsequent analyses. Finally, we kept 70 genomes containing 183 plasmids and 7 genomes with no plasmid according to the NCBI database (Supplementary table 2).

We filtered the assemblies based on contigs length (> 500 bp) and SPAdes coverage (> or = 2). Then, each assembly was mapped against the corresponding complete chromosome and plasmid sequences from the NCBI database using Quast 4.6 with standard parameters (13). Contigs that did not aligned on any sequence (chromosome and plasmid) or aligned on less than 50% of their length were not considered, as well as contigs that aligned on both sequences.

### PlaScope, Plasflow and cBar benchmark

PlaScope, Plasflow (5) and cBar (7) softwares were run on the reference dataset of 70 genomes containing plasmids. All these methods use different databases and classification approaches to sort contigs as plasmidic or chromosomal. Moreover, PlaScope and Plasflow may assign contigs as unclassified for ambiguous results.

For each tool, the predictions were considered as i) true positive (TP) (plasmid assignment of a plasmidic sequence), ii) true negative (TN) (chromosome or unclassified assignment of a non plasmidic sequence), iii) false positive (FP) (plasmid assignment of a non plasmidic sequence), iv) false negative (FN) (chromosome or unclassified assignement of a plasmidic sequence). Then, we calculated recall (TP/(TP+FN)), precision (TP/(TP+FP)), specificity (TN/(FP+TN)), accuracy ((TP+TN)/(TP+TN+FP+FN)) and F1 score (2*(recall*precision)/(recall+precision)). The results are presented for genomes taken as a whole in Table 1 and individually in Fig. 2.

**Figure 2.**
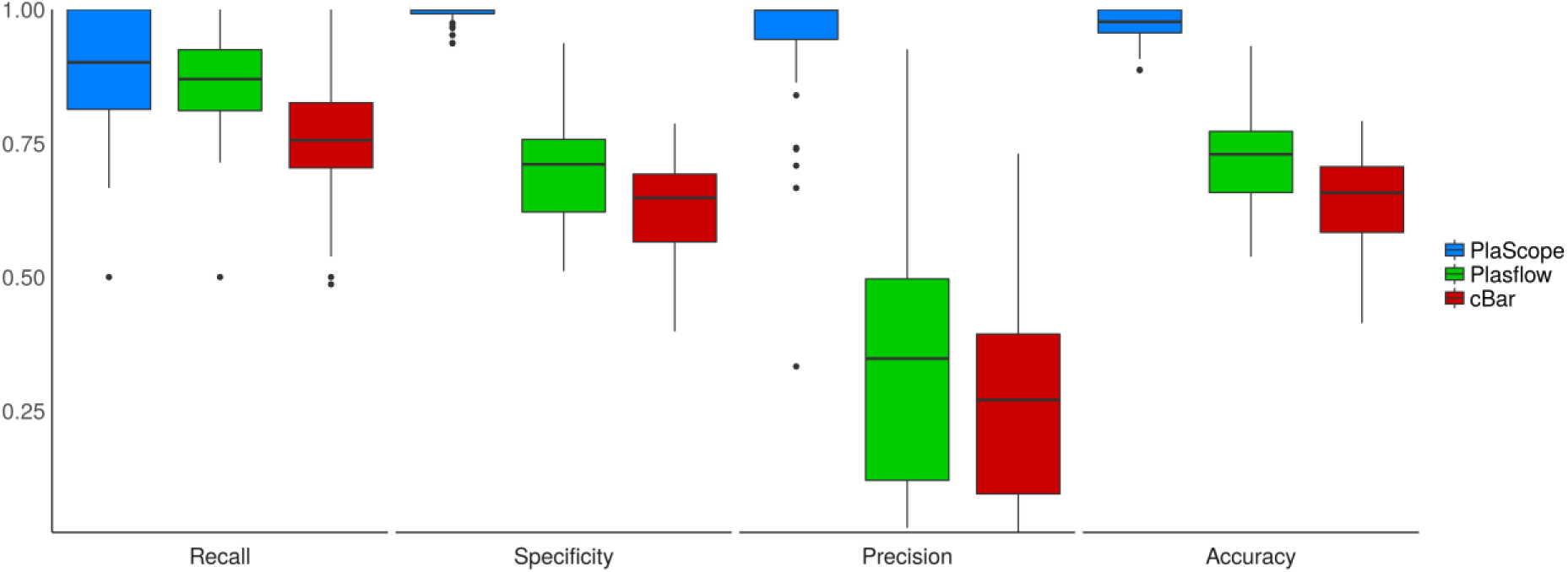
PlaScope, Plasflow and cBar performance for each plasmid taken individually. Recall, specificity, precision and accuracy obtained for each of the 183 plasmids are plotted according to the method in blue, green and red for PlaScope, Plasflow and cBar, respectively.

PlaScope achieves the highest recall on the dataset (0.87), closely followed by Plasflow (0.85), cBar being reaching the lowest value (0.74). However, important differences were found for the other assessment criteria. Indeed, we obtained with PlaScope very high precision (0.96), specificity (0.99) and accuracy (0.98) compared to Plasflow (0.27, 0.68 and 0.70, respectively) and cBar (0.21, 0.60 and 0.62, respectively). Obviously, these results are easily explained by the contents of our database, which was built specially for *E. coli*. Plasflow and cBar performed well in terms of recall, and their strength rely on their capacity to class many diverse taxonomic groups. Such methods can be really useful when working on metagenomes, but when focusing on a particular species a targeted approach like PlaScope drastically limits classification errors.

In addition, PlaScope was run on the 7 finished genomes with no plasmids. As expected, no plasmid was predicted for 6 genomes but, surprisingly, PlaScope predicted two plasmid contigs for *E. coli* KLY (GCA_000725305.1). To assess this result, we aligned these contigs against the NCBI database by blastN and obtained perfect alignments with the plasmid F sequence of *E. coli* K-12 C3026. This result suggests that the original assembly of *E. coli* KLY is missing this plasmid.

### Application to resistance, virulence gene and operon locations

In a second step, we evaluated our method on Extended-Spectrum Beta-lactamase (ESBL) carrying *E. coli* strains sequenced by Falgenhauer *et al.* (14). This dataset is particularly challenging because of an unusual high rate of chromosomal integration of CTX-M-15 coding genes. Indeed among the 27 isolates of sequence type (ST) 410, 21 carried a *blaCTX-M-15* gene on their chromosome. We downloaded short reads of these isolates and run PlaScope to classify the assembled contigs. In parallel, we determined the presence of CTX-M coding genes on the contigs using Resfinder (with a minimal identity of 95% and a minimal alignment coverage of 90%).

Using this approach, we accurately identified 20 chromosomally-integrated and 5 plasmid-related CTX-M (Fig. 3) compared to the publication results. We only had a discrepancy with the two isolates of Clade E (RS254 and RS371 strains). Indeed, we found a plasmid location of the CTX-M coding gene in the strain RS254 whereas it was described as chromosome-related, probably because of an uncommon structure formed by the gene and its adjacent sequences. For the second strain, RS371, the location was not predicted by PlaScope (unclassified) whereas it was stated as plasmid-located. The really short length of the contig carrying the CTX-M gene in this strain (i.e. 3274 bp) is certainly involved in this undetermined result.

**Figure 3.**
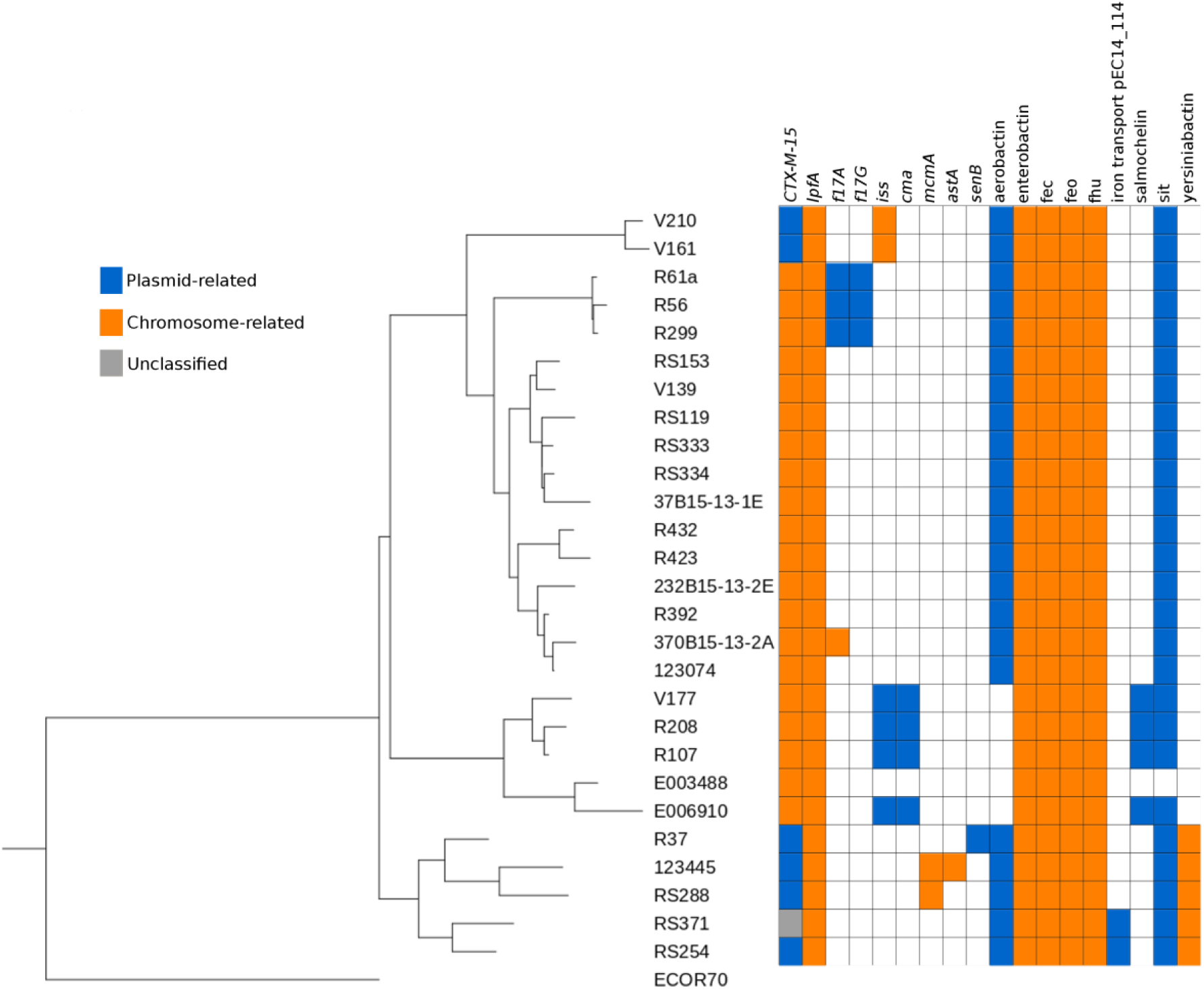
Genetic distance-based tree with location of *CTX-M-15*, virulence genes and operons in the ST410 *E. coli* strains from Falgenhauer *et al.* (14). We performed genome distance estimation with Mash (25) between all the strains and ECOR70 strain (26) used as an outgroup and known to be very close to ST410. Then we constructed a neighbour joining tree based on distance matrix with the module Phylo from biopython 1.68 (27), and generated an annotated tree with Interactive Tree Of Life (28). Location of the genes are displayed with colored squares (blue: plasmid prediction, orange: chromosome prediction, grey: unclassified).

In the same publication, the authors also searched for virulence genes and iron metabolism operons. To go further, we used PlaScope results to determine the location of these genes (Fig. 3). Some of them are exclusively carried by chromosomes (*lpfA*, *mcmA*, *astA*) or plasmids (*f17G*, *cma*, *senB*). Interestingly, *iss* can be found on either type of replicon. For example, *iss* is on chromosome in Clade A (V161 and V210 strains) isolates whereas it is located on plasmids in 4 out of the 5 Clade C (E003488, E006910, R107, R208 and V177 strains). This illustrates the different genetic background even between closely related strains. In the same way, the gene *f17A* has different locations: on plasmids in 3 strains (R299, R56, R61a) and on chromosome in only one (370B15-13-2A, not described in the original publication). These two possible locations of *iss* and *f17A* were previously observed (15, 16). Concerning the operons, 5 of them (i.e. enterobactin, fec, feo, fhu and yersiniabactin operons) were predicted as chromosome-related whereas the others, (i.e. aerobactin, salmochellin, sit and the iron transport pEC14_114) were predicted as plasmidic. These results are in agreement with the literature. Indeed, the first five are known to be chromosome-encoded (17-21) whereas iron transport pEC14_114 is plasmidic (22). Aerobactin, salmochelin and sit have been found on both types of replicons (23).

## Conclusion

Here, we propose a method, called PlaScope, for plasmid and chromosome classification of *E. coli* contigs. It is based on Centrifuge (10): a fast metagenomic classifier that uses exact matches and small-sized databases. PlaScope offers a high specificity by selecting a unique assignment of contigs to plasmid, chromosome or unclassified. Indeed, we took advantage of the ever growing number of sequences from databases to build a custom database, which combines many high quality sequences of *Enterobacteriaceae* plasmids and chromosome sequences of *E. coli*. We compared the performance of our tool with cBar and Plasflow, as these bioinformatic softwares also enable the segregation of plasmid and chromosome contigs. These two programs rely on genomic signature and have been develop to predict plasmid sequences in metagenomic samples.

Compared to PlaScope, Plasflow achieve roughly the same recall value on our dataset, whereas cBar performed a little bit less well. However when looking at the other criteria such as precision, specificity and accuracy, PlaScope outperformed the other ones due to its highly specific database. cBar and Plasflow are virtually able to identify mobile elements in many bacterial species owing to their very diverse taxonomic database. But when focusing on a species, the targeted approach of PlaScope gave indisputably better results both in terms of recall and precision.

Using PlaScope, we were able to recover almost all plasmids from the analysed strains, with very high precision, specificity and accuracy. Furthermore, among 1 of the 7 strains described as non-bearing plasmid strains in the NCBI database we were able to identify a mobile element: a typical plasmid F in a *E. coli* K-12.

In a second analysis, we challenged our approach on more concrete data by looking at specific genes. Analysing clinical or environmental strains, it could be of great interest to detect specific clones with particular genetic backgrounds. Indeed, plasmid location of a resistance or virulence gene has not the same impact on an epidemiological point of view and on the capacity of transmission of the strain in a particular environment. For example, we can fear plasmid outbreaks when a gene that confers resistance against wide-spectrum antibiotic is carried by such a mobile element. Conversely, if the same gene integrates in the chromosome of an already highly virulent strain, it can lead to the emergence of a well-adapted and dangerous clone. To highlight this, we chose a genome dataset of *E. coli* wherein many strains exhibited a chromosomal integration of CTX-M-15 coding gene, one of the main enzymes responsible for resistance to wide spectrum antibiotics such as cephalosporins in *E. coli* (14). Using PlaScope we accurately identify 20/21 of these chromosomal insertions. Beside, we predicted the location of virulence genes and iron metabolism operons and it was in agreement with the literature. It demonstrates that PlaScope may be really useful to locate operons like aerobactin or salmochellin, which can be on plasmid as well as chromosome and have, like other iron-metabolism related systems, major impact on virulence and/or fitness (21, 24).

We think that our approach can be very useful when focusing on a well-described species as it makes it possible to decipher the plasmid content of the genomes without an excess of over prediction. It can highlight integration events or plasmid transmission between isolates. Nonetheless as it is based on previous knowledge of plasmids found in a specific taxon *(e.g. Enterobacteriaceae*), it will require the enrichment of the database keeping it up-to-date. At last, it could also be interesting to create other databases for well-known bacteria with many complete genomes available such as *Klebsiella pneumoniae, Staphylococcus aureus* or *Bacillus* species.

